# Structural basis of TFIIH activation for nucleotide excision repair

**DOI:** 10.1101/628032

**Authors:** Goran Kokic, Aleksandar Chernev, Dimitry Tegunov, Christian Dienemann, Henning Urlaub, Patrick Cramer

**Author notes:** Correspondence and requests for materials should be addressed to PC.

## Abstract

Genomes are constantly threatened by DNA damage, but cells can remove a large variety of DNA lesions by nucleotide excision repair (NER)^1^. Mutations in NER factors compromise cellular fitness and cause human diseases such as Xeroderma pigmentosum (XP), Cockayne syndrome and trichothiodystrophy^2,3^. The NER machinery is built around the multisubunit transcription factor IIH (TFIIH), which opens the DNA repair bubble, scans for the lesion, and coordinates excision of the damaged DNA single strand fragment^1,4^. TFIIH consists of a kinase module and a core module that contains the ATPases XPB and XPD^5^. Here we prepare recombinant human TFIIH and show that XPB and XPD are stimulated by the additional NER factors XPA and XPG, respectively. We then determine the cryo-electron microscopy structure of the human core TFIIH-XPA-DNA complex at 3.6 Å resolution. The structure represents the lesion-scanning intermediate on the NER pathway and rationalizes the distinct phenotypes of disease mutations. It reveals that XPB and XPD bind double- and single-stranded DNA, respectively, consistent with their translocase and helicase activities. XPA forms a bridge between XPB and XPD, and retains the DNA at the 5’-edge of the repair bubble. Biochemical data and comparisons with prior structures^6,7^ explain how XPA and XPG can switch TFIIH from a transcription factor to a DNA repair factor. During transcription, the kinase module inhibits the repair helicase XPD^8^. For DNA repair, XPA dramatically rearranges the core TFIIH structure, which reorients the ATPases, releases the kinase module and displaces a ‘plug’ element from the DNA-binding pore in XPD. This enables XPD to move by ~80 Å, engage with DNA, and scan for the lesion in a XPG-stimulated manner. Our results provide the basis for a detailed mechanistic analysis of the NER mechanism.

---

A mechanistic dissection of the NER machinery was thus far hampered because TFIIH was not available in large quantities. We therefore established protocols to prepare milligram amounts of recombinant human TFIIH core and kinase modules (Methods). The purified TFIIH core comprised seven subunits including the ATPases XPB and XPD, whereas the kinase module contained CDK7, cyclin H, and MAT1. From these two modules we could reconstitute the complete 10-subunit TFIIH (Extended Data Fig. 1a).

To analyse the enzymatic activities of TFIIH, we monitored helicase and translocase activities in real time by fluorescence resonance energy transfer (Methods). Core TFIIH showed 5’-3’ helicase activity, which was lost upon mutation of the XPD active site (Fig. 1a, Extended Data Fig. 1b). This is consistent with a prior description of XPD as a 5’-3’ DNA helicase^8,9^. Core TFIIH also showed translocase activity, which was however much lower than the helicase activity (Fig. 1b). Translocase activity was due to XPB because it was sensitive to triptolide (Extended Data Fig. 1c), a drug that targets XPB^10^. This is consistent with the known translocase activity for the yeast XPB homologue^11^, but not with helicase activity reported for an archaeal XPB homologue^12^.

**Figure 1.**
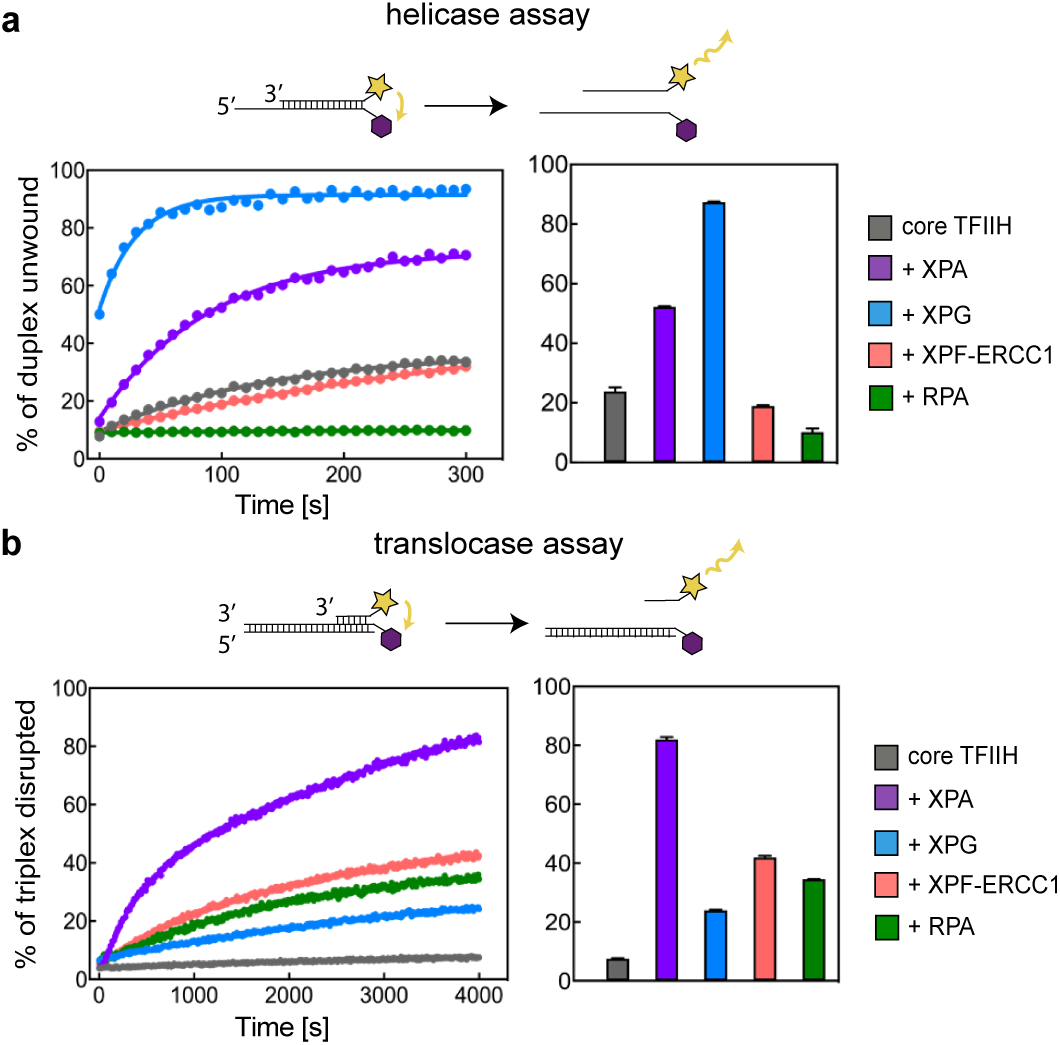
Regulation of ATPases in core TFIIH. **a.** Effect of NER proteins on XPD 5’-3’ helicase activity. Real-time fluorescence measurement using a fluorescence energy transfer-based assay. Bars show percentage of unwound product after 100s. Error bars provide standard deviations of the mean of two replicates. RPA inhibits XPD helicase activity, by masking the single-stranded DNA overhang. **b.** Effect of NER proteins on XPB translocation activity. Real-time fluorescence measurement of triplex disruption. Bars show percentage of disrupted triplex after 4000s. Error bars provide standard deviations of the mean of two replicates.

To test whether other NER factors affect TFIIH activities, we purified human XPA, XPG, RPA, and XPF-ERCC1 complex (Extended Data Fig. 1d). In the presence of XPA or XPG, DNA unwinding by XPD was four-fold or 20-fold faster, respectively, as deduced from stopped-flow kinetics (Fig. 1a, Extended Data Fig. 1e). A stimulation of XPD by XPA was observed before^9,13^, but the effect of XPG on DNA unwinding is much stronger. This explains the earlier observation that XPG is required for efficient DNA bubble opening^14^ and implicates XPG in lesion scanning by XPD. XPB translocation activity was stimulated by all NER factors tested, although stimulation by XPA was exceptionally strong (Fig. 1b).

We now investigated the structural basis for how XPA and XPG activate the TFIIH ATPases. We prepared the core TFIIH-XPA-XPG complex bound to a bifurcated DNA scaffold that mimics one half of a DNA repair bubble (Extended Data Fig. 2a). We imaged this complex by cryo-EM and solved the structure at an overall resolution of 3.6 Å (Methods, Extended Data Figs. 2, 3). The cryo-EM density was of high quality and revealed DNA and all protein components except XPG, which likely dissociated during cryo-EM grid preparation. The derived structure contains the p52 subunit and other regions that were lacking from the previous human TFIIH structure^6^ and reveals the XPB-TFIIH core interface (Fig. 2, Extended Data Fig. 3f).

**Figure 2.**
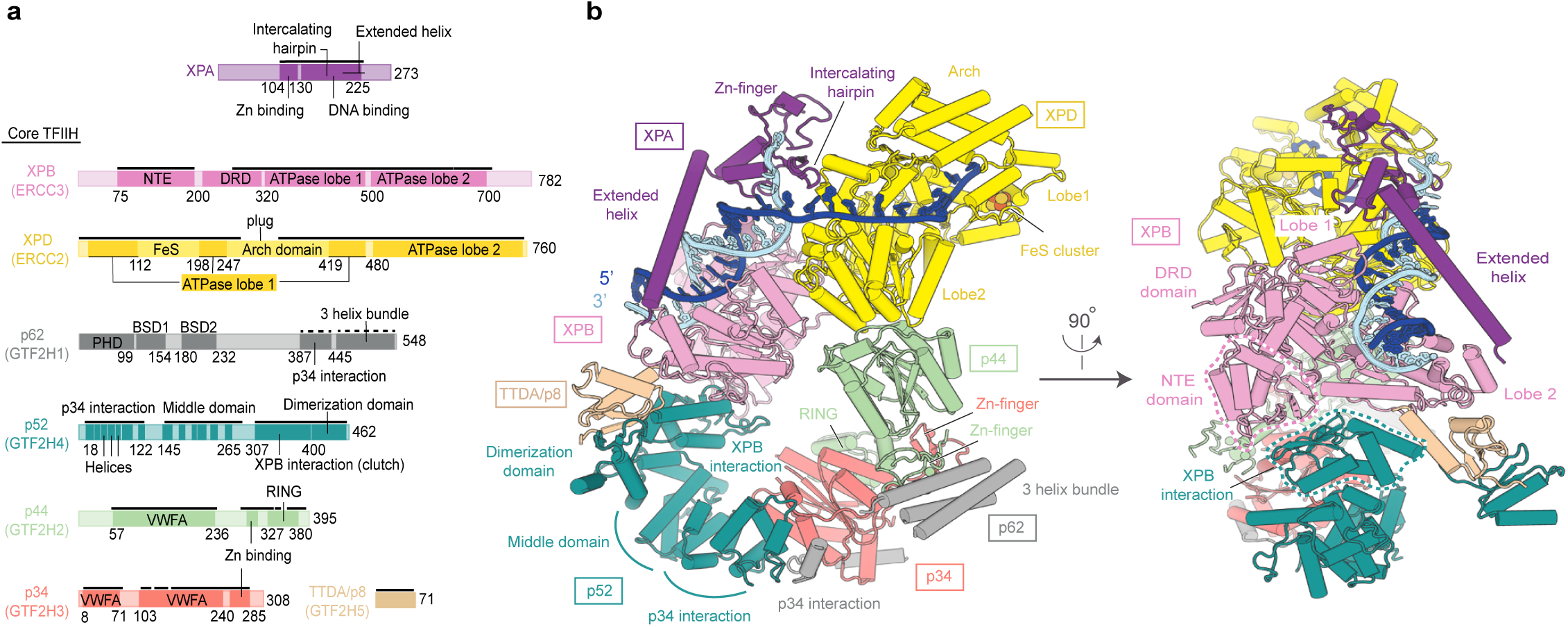
Structure of human core TFIIH-XPA-DNA complex. **a.** Domain organization of XPA and human TFIIH subunits. Residues at domain borders are indicated. Solid and dashed black lines mark residues modeled as atomic and backbone structures, respectively. DRD, damage recognition domain; NTE, N-terminal extension; BSD, BTF2-like, synapse-associated and DOS2-like domains; VWFA, von Willebrand factor type A domain. **b.** Cylindrical representation of the structure. Proteins colored as in (a). Newly build XPB and p52 domains are highlighted with dashed lines and reveal how is XPB connected to the TFIIH core.

The structure of the core TFIIH-XPA-DNA complex differs substantially from the TFIIH structure observed in transcription complexes (Fig. 2)^7,15^. Both XPB and XPD bind DNA (Extended Data Fig. 4), whereas only XPB binds DNA in transcription complexes^7,15^. XPB binds DNA in the duplex region, whereas XPD binds the single-stranded 3’-DNA extension, consistent with translocase and helicase function, respectively. DNA binding of both ATPases requires large structural changes in TFIIH (Supplementary Video 1). XPD and its associated subunit p44 move by ~80 Å, and this requires a flexible connection between subunits p44 and p34, and rearrangements in subunit p52 (Extended Data Fig. 5).

The structure informs on XPA, which is essential for NER^16^. XPA contains an N-terminal zinc finger and a DNA-binding domain with an extended helix and an intercalating ß-hairpin (Fig. 2). XPA forms an elongated arch over DNA that bridges between the two ATPases. XPA binds XPB and XPD with its extended helix and its intercalating hairpin, respectively. The C-terminal region of XPA extends to p52 and TTDA/p8 (Extended Data Figs. 3d, 7c), explaining why TTDA/p8 facilitates XPA recruitment *in vivo*^17^.

These observations explain how XPA stimulates XPB translocation. First, XPA connects both XPB ATPase lobes to p52 and TTDA/p8 subunits that stimulate the XPB activity within the TFIIH core^18,19^. Second, XPB is not a processive enzyme and readily dissociates from DNA^11^. However, in our structure DNA is held in a positively charged DNA duplex tunnel that is formed between the extended helix of XPA and the XPB ATPase (Fig. 3a). XPA thus retains DNA near the XPB active site. Indeed, our cryo-EM data revealed an alternative state of the complex with the DNA duplex disengaged from XPB but retained by the XPA extended helix (Fig. 3b, Extended Data Fig. 2d). Thus, by trapping the DNA within the duplex tunnel, XPA may retain the NER machinery on the DNA during lesion scanning and processing.

**Figure 3.**
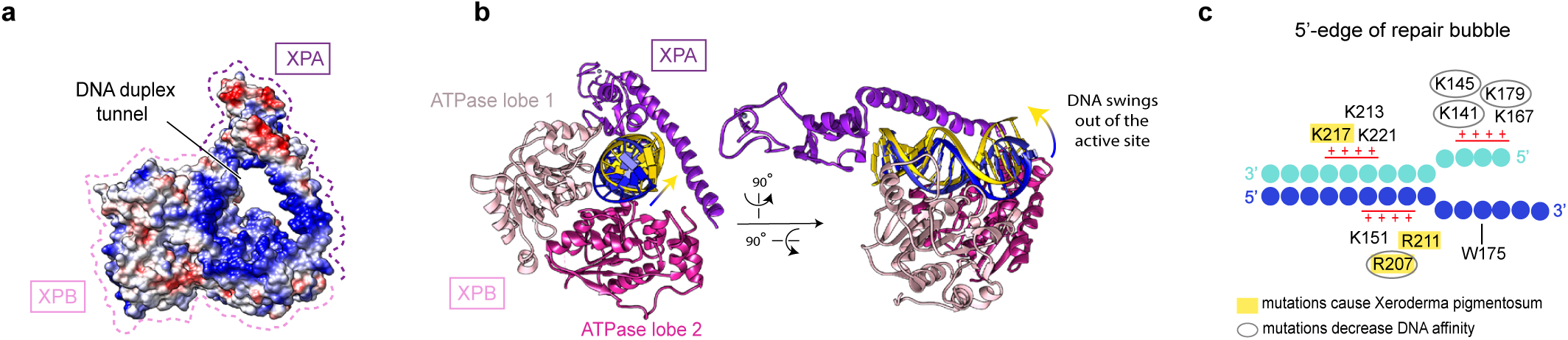
XPA-DNA interactions. **a.** DNA duplex tunnel formed by XPA and XPB. Blue, white and red color indicates positive, neutral and negative electrostatic surface potential, respectively. Created with UCSF Chimera^35^. **b.** Two positions of DNA in the tunnel. Tightly bound DNA is in blue, dissociated DNA in yellow, ATPase lobe 1 of XPB in pink, ATPase lobe 2 in hotpink, and XPA in purple. **c.** Electrostatic interactions between XPA and the DNA junction. DNA nucleotides are indicated as circles. Patches of positively charged residues in proximity to the DNA backbone are indicated. Residues that are mutated in Xeroderma pigmentosum^16^ are highlighted in yellow. Mutation of encircled residues decreases DNA affinity^36^.

XPA also contributes to the recognition of the 5’-edge of the DNA repair bubble that depends on electrostatic interactions (Fig. 3c). XPA inserts its intercalating ß-hairpin between DNA single strands at the duplex junction (Fig. 2b), consistent with published biochemical data^20^. The tip of the XPA hairpin contains a conserved tryptophan residue (Trp175) that stacks against the base of the DNA 3’-extension at the junction (Extended Data Fig. 3i). Several sites of mutations that cause severe *Xeroderma pigmentosum*^16^ map to XPA residues that interact with DNA (Fig. 3c). Previous studies of the yeast XPA counterpart suggested that XPA may detect DNA lesions^21^. However, our results suggest that XPA rather demarcates the 5’-edge of the repair bubble and stimulates lesion scanning by clamping core TFIIH onto DNA.

The detailed structure also enables the localization of XPD residues mutated in patients with *Xeroderma pigmentosum* (XP) and *Trichothiodystrophy* (TTD) and rationalizes their functional effects, as previously suggested by biochemical studies^22^ and comparisons to archaeal XPD homologues in the absence of DNA^23-25^ (Extended Data Fig. 6, Supplementary Note 1). Most notably, the XP mutations affect the DNA-contacting residues (Extended Data Fig. 6b) in the ATPase lobe 2 which would specifically impair the XPD helicase activity and DNA repair. In contrast, TTD mutations map to the XPD-p44 interface, around the FeS cluster and between the XPD ATPase lobes (Extended Data Fig. 6c-e). Thus, TTD mutations would compromise the integrity of TFIIH and XPD stability, thereby affecting transcription and other TFIIH-mediated processes outside NER^5^, which leads to more severe phenotypes.

The structure also provides details of the XPD-DNA interactions (Fig. 4a, b). The ATPase lobe 2 interacts with DNA bases near the duplex-single strand junction, which includes base stacking with the side chains of residues F508 and Y627. This mode of DNA interaction is unusual for helicases of the SF2 family, which generally engage with the sugar-phosphate backbone^26,27^. We speculate that the extensive contacts of XPA and XPD with single-stranded DNA facilitate DNA opening and XPD loading during initial stages of NER^28^.

**Figure 4.**
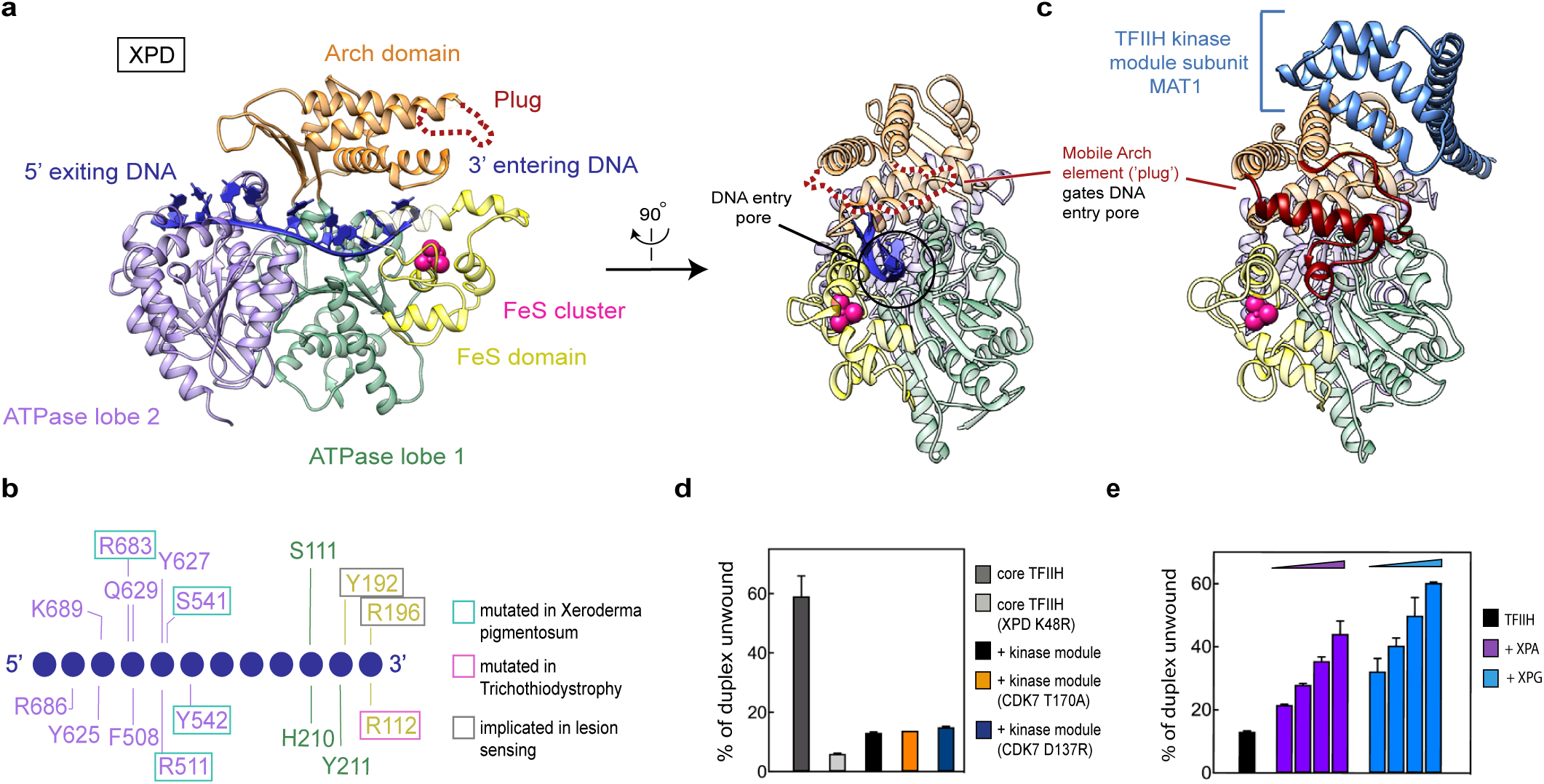
XPD activation and DNA binding. **a.** Two views of XPD bound to DNA. XPD domains ATPase lobe 1, FeS, Arch, and lobe 2 are in green, yellow, orange, and medium purple, respectively. DNA is dark blue. A black circle depicts the DNA pore. **b.** Schematic representation of XPD-DNA interactions. **c.** Side view of XPD bound to the kinase module (PDB code 5OF4)^6^. The plug in the Arch domain is in dark red, the kinase module subunit MAT1 in blue. **d.** Effect of kinase module variants on XPD helicase activity. Core TFIIH was incubated with 2-fold excess of the kinase module and helicase activity was monitored as in Figure 1a. Bars show the percentage of unwound product after 300s for two replicates. Error bars represent s.d. from the mean values. **e.** Effect of increasing concentrations of XPA and XPG on XPD helicase activity in the presence of kinase module (0.25 µM core TFIIH, 0.5 µM kinase module, 0.375, 0.75, 1.5 or 3 µM XPA or XPG). Otherwise as in (d).

The structure further suggests how XPD verifies the lesion during DNA scanning. The DNA single strand extends into a pore formed by the ATPase lobe 1, the iron-sulfur cluster (FeS) domain, and the Arch domain (Fig. 4a). The sugar-phosphate backbone is bound by residues in the FeS domain, including Y192 and R196, which were implicated in DNA lesion sensing^29^ (Fig. 4b). Residues R112 and C134 bridge between DNA and the FeS cluster (Extended Data Fig. 4d), which was suggested to be involved in lesion detection via DNA-mediated charge transfer^30^. The FeS cluster is flanked by two protein pockets that are lined with aromatic residues Y158, F161 and F193, and may proof-read DNA bases, as observed for base excision DNA repair^31^.

We could reproduce the known repression of the XPD helicase activity by the TFIIH kinase module^8,9^ in our helicase assays (Fig. 4d). We found that catalytically inactive variants of the TFIIH kinase module (Extended Data Fig. 1f) could also repress XPD helicase activity, showing that repression does not require the kinase activity of CDK7 (Fig. 4d). Both XPA and XPG could relieve the kinase-mediated repression of XPD in a concentration-dependent manner (Fig. 4e). These observations show that XPA and XPG counteract the repressive effect of the kinase module on XPD.

A comparison of our structure with the previous TFIIH structures shows how the kinase module represses XPD activity. In previous TFIIH structures^6,7^, a region in the XPD Arch domain forms a ‘plug’ (residues 273-325) that occupies the DNA pore of XPD (Fig. 4c). The plug would clash with DNA in the XPD pore, but is displaced and mobile in our structure (Fig. 4a). The kinase module subunit MAT1 contacts the plug and may stabilize it in the XPD pore, explaining how the kinase module impairs binding of core TFIIH to single stranded DNA and XPD helicase activity^9^. In addition, a loop in the yeast counterpart of p62 extends into the XPD active site^7^, and would interfere with the observed DNA trajectory through the helicase.

Structural comparisons also suggest how XPA relieves XPD inhibition by the kinase module. XPA stabilizes TFIIH in a new conformation in which the two ATPases are drastically reoriented. This conformation is incompatible with MAT1 binding as observed in the previous TFIIH structure^6^ (Extended Data Fig. 5e). This also explains how XPA facilitates kinase module removal upon NER induction *in vivo*^32^. Taken together, MAT1 and XPA stabilize two entirely different conformations of TFIIH, which contain the repair helicase XPD in an inactive or an active state, respectively.

Since XPG was not visible in our structure, we located it by chemical crosslinking (Extended Data Fig. 7). The crosslinking data matches the structural data very well (Extended Data Fig. 7c), and unambiguously localizes XPG. The N-terminal region of XPG specifically crosslinks to the FeS and Arch domains in XPD, including the plug element, whereas the C-terminal extension of XPG crosslinks mostly to XPB and p52 (Extended Data Fig. 7d). In addition, XPG crosslinks to XPD at its binding site for the kinase module (Extended Data Fig. 7d), suggesting that XPG competes with the kinase module for XPD binding, and explaining how XPD inhibition by the kinase module is relieved (Fig. 4e). These data suggest that XPG facilitates lesion scanning by blocking the kinase module-binding site on XPD and directly stimulating XPD helicase activity. We note that XPG may bind to TFIIH in alternative ways during other TFIIH functions, in which XPD is not bound to DNA^5-7,33^.

Taken together, our structure-function analysis extends our understanding of the NER pathway (Extended Data Fig. 8). The DNA lesion is first recognized by XPC, which recruits TFIIH^1,13^. XPA and XPG then displace the kinase module, stabilize an alternative conformation of TFIIH, and remove the inhibitory plug from the XPD pore. XPA and XPG also stimulate XPB and XPD, and this may facilitate DNA opening^4,14,18,19^ and XPD migration in the 3’ direction to scan the DNA strand for the lesion^13^. XPA chaperons TFIIH-DNA interactions and anchors the NER machinery to the 5’ edge of the repair bubble, where it is ideally positioned to recruit XPF-ERCC1 and complete the repair assembly when the lesion is encountered^16^. The two endonucleases XPF and XPG can now incise the lesion-containing DNA strand near the 5’- and the 3’-edge of the repair bubble, respectively, to remove the lesion-containing DNA fragment^1^. We acknowledge that while our manuscript was in revision, a manuscript became available online^34^ that provides a high resolution free TFIIH structure and includes a mapping of disease mutations consistent with the one described here.

## Acknowledgements

We thank current and former members of Cramer Laboratory, including C. Bernecky, C. Burzinski, S. Dodonova, H. Hillen, S. Vos and F. Fischer. GK was supported by a Boehringer-Ingelheim PhD Fellowship. GK and AC have been doctoral students of the Ph.D. program “Molecular Biology” – International Max Planck Research School and the Göttingen Graduate School for Neurosciences, Biophysics, and Molecular Biosciences (GGNB) (DFG grant GSC 226) at the Georg August University Göttingen. PC was supported by the Deutsche Forschungsgemeinschaft (SFB860, SPP1935), the European Research Council Advanced Investigator Grant TRANSREGULON (grant agreement No 693023), and the Volkswagen Foundation.

## Author contributions

GK designed and carried out all experiments except for crosslinking-mass spectrometry, which was carried out by AC. DT assisted with image processing and CD assisted with cryo-EM data acquisition and model building. HU supervised mass spectrometry. PC designed and supervised research. GK and PC interpreted the data and wrote the manuscript, with input from all authors.

## Author information

The authors declare no competing financial interest.

## METHODS

### Cloning and protein expression

Vectors encoding full-length XPA, XPG, XPF, ERCC1, RPA1, RPA2, RPA3, XPB, XPD, p62, p52, p44, p34, MAT1, CDK7, cyclin H and TTDA were ordered from Harvard Medical School PlasmID Repository and used as a template for gene amplification by PCR. Amplified genes were cloned into MacroBac vectors via ligation independent cloning, as described^37^. ERCC1, p52, p34 and TTDA were cloned into 438A (Addgene: 55218), XPA, XPF, XPB, p62, p44, MAT1, CDK7 and CycH into 438B (Addgene: 55219) and XPG and XPD into 438C vector (Addgene: 55220) which resulted in no tag, N-terminal 6x His or N-terminal 6x His followed by a maltose binding protein, respectively. XPF and ERCC1, MAT1, CDK7 and CycH, as well as XPB, p62, p52, p44, p34 and TTDA were combined into a single vector via restriction digestion and ligation independent cloning^37^. RPA1 was cloned in 11B (Addgene: 48295), and RPA 2 and 3 were cloned into 11A vectors (Addgene: 48294), followed by assembly of all RPA subunits into one vector by successive ligation independent cloning reactions. All tags were separated from the gene with a tobacco etch virus protease site. All mutant proteins used here (CDK7:D137R, CDK7:T170A, XPG:E791A, XPD:K48R) were produced by round-the-horn side-directed mutagenesis and purified as their wild type counterparts.

RPA complex was expressed in *E. coli* and purified as described^38^. TFIIH kinase module comprised of MAT1, CDK7 or CDK7 mutants and cyclin H was expressed and purified as described^39^. All other proteins and protein complexes were expressed in insect cells and purified as described below. Bacmid preparation, V0 and V1 virus production was described previously^40^. Core TFIIH was produced by co-infecting the cells with two V1 viruses: virus encoding XPB, p62, p52, p44, p34 and TTDA and a virus encoding XPD or XPD mutant. All proteins were expressed in Hi5 cells grown in ESF-921 media (Expression Systems, Davis, CA, United States) at 27 °C. Typically, 600 ml culture was infected with 500 μL of V1 virus and grown for 48 – 72 hrs prior harvesting by centrifugation (30 min, 4°C, 500 xg). Cell pellet was resuspended in lysis buffer; 400 mM KCl, 20 % glycerol (v/v), 20 mM KOH-HEPES pH 7, 5 mM β-mercaptoethanol, 30 mM imidazole pH 8.0, 0.284 μg/ml leupeptin, 1.37 μg/ml pepstatin A, 0.17 mg/ml PMSF and 0.33 mg/ml benzamidine for the core TFIIH, and 400 mM NaCl, 20 mM Tris-HCl pH 7.9, 10 % glycerol (v/v), 1 mM DTT, 30 mM imidazole pH 8.0, 0.284 μg/ml leupeptin, 1.37 μg/ml pepstatin A, 0.17 mg/ml PMSF and 0.33 mg/ml benzamidine for other proteins. Cell suspension was flash frozen in liquid nitrogen and stored at −80 °C.

### Protein purification

All purification steps were performed at 4 °C and all buffers were filtered and thoroughly degassed immediately before use. Cells were thawed in a water bath operating at 30 °C and opened by sonication. The lysate was clarified by centrifugation (18,000 xg, 30 min), followed by ultracentrifugation (235,000 xg, 60 min). In case of core TFIIH the clarified lysate was first filtered using 0.8 µm syringe filters (Millipore) and loaded onto HisTrap HP 5 mL (GE Healthcare, Little Chalfont, United Kingdom). The column was washed with 10 CV of lysis buffer, followed by 20 CV of high salt wash (800 mM KCl, 20 % glycerol (v/v), 20 mM KOH-HEPES pH 7, 5 mM β-mercaptoethanol, 30 mM imidazole pH 8.0, 0.284 μg/ml leupeptin, 1.37 μg/ml pepstatin A, 0.17 mg/ml PMSF and 0.33 mg/ml benzamidine). Column was washed again with 5 CV of lysis buffer and protein was subsequently eluted with a gradient of 0–80% elution buffer (400 mM KCl, 20 % glycerol (v/v), 20 mM KOH-HEPES pH 7, 5 mM β-mercaptoethanol, 500 mM imidazole pH 8.0, 0.284 μg/ml leupeptin, 1.37 μg/ml pepstatin A, 0.17 mg/ml PMSF and 0.33 mg/ml benzamidine). Fractions were checked on NuPAGE 4-12 % Bis – Tris Protein Gels (Invitrogen) for the presence of all core TFIIH subunits and appropriate fractions were pulled and mixed with 10 ml of amylose resin (New England BioLabs) pre-equilibrated in washing buffer (400 mM KCl, 20 % glycerol (v/v), 20 mM KOH-HEPES pH 7, 5 mM β-mercaptoethanol, 2 mM MgCl_2_ and 10 μM ZnCl_2_). The protein solution was incubated with the beads for 1h while rotating. The amylose resin was poured into Econo-Pac Chromatography columns (BioRad) and washed with 5 CV of washing buffer. Protein was eluted with washing buffer containing 100 mM maltose. Protein-containing fractions were pooled, mixed with 2 mg of TEV protease and dialysed against 2 L of dialysis buffer overnight (250 mM KCl, 20 % glycerol (v/v), 20 mM KOH-HEPES pH 7, 5 mM β-mercaptoethanol, 2 mM MgCl_2_ and 10 μM ZnCl_2_). The dialysed sample was applied to DEAE (GE Healthcare) and heparin column (GE Healthcare) in tandem and washed with 20 CV of dialysis buffer. After the removal of DEAE column, protein was eluted with a gradient of elution buffer 0-100% (1M KCl, 20 % glycerol (v/v), 20 mM KOH-HEPES pH 7, 5 mM β-mercaptoethanol, 2 mM MgCl_2_ and 10 μM ZnCl_2_). Peak fractions were pooled, concentrated with Amicon Millipore 15 ml 100,000 MWCO centrifugal concentrator and applied to Superose 6 increase 10/300 GL column (GE Healthcare) equilibrated in storage buffer (400M KCl, 20 % glycerol (v/v), 20 mM KOH-HEPES pH 7.0, 5 mM β-mercaptoethanol, 2 mM MgCl_2_). Peak fractions were again concentrated, aliquoted, flash frozen and stored at −80 °C.

XPF-ERCC1, XPG and XPA containing lysate was applied onto GE HisTrap HP 5 mL (GE Healthcare, Little Chalfont, United Kingdom) equilibrated in lysis buffer (in case of XPA all downstream steps were performed in the presence of 5 mM β-mercaptoethanol and 10 μM ZnCl_2_ instead of 1 mM DTT). The column was washed with 20 CV of high salt buffer (800 mM NaCl, 20 mM Tris-HCl pH 7.9, 10 % glycerol (v/v), 1 mM DTT, 30 mM imidazole pH 8.0, 0.284 μg/ml leupeptin, 1.37 μg/ml pepstatin A, 0.17 mg/ml PMSF and 0.33 mg/ml benzamidine). After 5 CV wash with the lysis buffer, proteins were eluted with the elution buffer gradient 0-80% (400 mM NaCl, 20 mM Tris-HCl pH 7.9, 10 % glycerol (v/v), 1 mM DTT, 500 mM imidazole pH 8.0, 0.284 μg/ml leupeptin, 1.37 μg/ml pepstatin A, 0.17 mg/ml PMSF and 0.33 mg/ml benzamidine). Pulled peak fractions were processed differently for different proteins. XPF-ERCC1 and XPA containing protein solutions were directly mixed with 2 mg of TEV protease and dialysed against 1L of dialysis buffer (400 mM NaCl, 20 mM Tris-HCl pH 7.9, 10 % glycerol (v/v), 1 mM DTT). XPG solution was mixed with 10 ml of amylose resin (New England BioLabs) pre-equilibrated in dialysis buffer. Solution was mixed for 1 hr and subsequently poured into Econo-Pac Chromatography columns (BioRad). Column was washed with 10 CV of dialysis buffer followed by elution with dialysis buffer containing 100 mM maltose. Protein containing fractions were pulled, mixed with 2 mg of TEV protease and dialysed against 1L of dialysis buffer. Dialysed solutions containing XPF:ERCC1, XPA and XPG were loaded on GE HisTrap HP 5 mL and flow through fractions were collected. Protein containing fractions were checked for contaminants on NuPAGE 4-12 % Bis – Tris Protein Gels (Invitrogen), pulled, concentrated with appropriate Amicon Millipore 15 ml centrifugal concentrator and applied onto Superdex 75 10/300 equilibrated in storage buffer (400 mM NaCl, 20 mM NaOH:HEPES pH 7.5, 10% glycerol (v/v), 10 μM ZnCl_2_ and 5 mM β-mercaptoethanol) for XPA and Superdex 200 10/300 increase (GE Healthcare) equilibrated in a different storage buffer (400 mM NaCl, 20 mM NaOH:HEPES pH 7.5, 10% glycerol (v/v), 1mM DTT) for the rest of the proteins. Peak fractions were pooled, concentrated, aliquoted, flash frozen and stored at −80 °C.

### Helicase and translocase assays

H1 and H2 DNA sequences (Extended Data Table 1) were used for monitoring the helicase activity in 5’-3’ direction and H3 and H4 for monitoring the helicase activity in 3’-5’ direction. DNA annealing reaction contained fluorescent DNA primer (25 μM) and quenching DNA oligo (37.5 μM) dissolved in water. Annealing was performed in a thermocycler by heating up the DNA solution to 95 °C for 5 min, followed by slow cooling (1 °C / min) to 4 °C. Typical unwinding reactions of 20 μl final volume contained 0.4 pmol of DNA duplex and 8 pmol of core TFIIH in 100 mM KCl, 20 mM KOH:HEPES pH 7, 5% glycerol, 0.2 mg/ml BSA, 3 mM phosphoenolpyruvate, 10 mM MgCl2, 1 mM DTT and excess amount of pyruvate kinase (Sigma). When the effect of DNA repair factors on unwinding was measured we supplemented the reaction with 24 pmol of the corresponding factor. The reaction mixture was preincubated at 26 °C for 10 min. The reaction was started by addition of ATP (2 mM final) and the unwinding was monitored at 26 °C by using the Infinite M1000Pro reader with excitation wavelength 495 nm, emission wavelength 520 nm and gain of 150. Percentage of unwound product was calculated by dividing the observed fluorescence intensity by the intensity of the fluorescent primer in the reaction buffer (mimicking fully unwound DNA).

DNA unwinding monitored by stopped-flow was performed in the same buffer conditions and with the same final protein and DNA concentrations as above. The core TFIIH preincubated with XPA or XPG was rapidly mixed with equal volume of ATP (2mM final) in the SX-20MV stopped-flow apparatus (Applied Photophysics). FAM fluorescence was monitored upon excitation at 465 nm after passing through KV500 cut-off filter (Schott). All time courses shown represent average of 5 technical replicates. Initial rate of DNA unwinding was calculated using Prism (Graphpad software) by fitting the initial linear part of the fluorescence trace.

Triplex displacement assay was performed in a similar way as previously described^11^. 10 μl annealing reaction for triplex displacement assay contained T1 (30 μM) and T2 (25 μM) oligo (Extended Data Table 1) in 25 mM MES pH 5.5 and 10 mM MgCl_2_. The reaction was heated to 95 °C for 5 min followed by slow cooling (1 °C/min) to 4 °C. After cooling, the reaction was supplemented with 1 μl of florescent T3 oligo (9 μM final), heated to 57 °C and cooled down to 20 °C at the speed of 1 °C/min. Translocation reactions were preformed exactly as described for the helicase assay, only with triplex DNA as a substrate. A higher core TFIIH input (75 pmol in 20 μl reactions) was used when no stimulatory factors were added (Extended Data Fig. 1c) to obtain a more robust fluorescent signal.

### Kinase activity assay

We used the kinase activity assay to assess the activity of kinase module variants containing CDK7:D137R or CDK7:T170A mutants^41^. As CDK7 phosphorylates the C-terminal domain of the largest RNA polymerase II subunit during transcription initiation, we used purified yeast RNA polymerase II^42^ dephosphorylated with lambda phosphatase during purification^39^ as a substrate in the assay. RNA polymerase II (50 nM final) was mixed with increasing concentrations of kinase module variants (30, 100, 220 and 500 nM final) in final buffer conditions containing 100 mM KCl, 20 mM KOH:HEPES pH 7.5, 3 mM MgCl_2_, 5% glycerol and 5 mM β-mercaptoethanol, and preincubated for 2 min at 30 °C. Reactions were started by the addition of ATP (0.5 mM final) and quenched after 2 min at 30 °C with EDTA (100 mM) and 4X LDS buffer (Invitrogen). Reactions were run on 4-12% Bis-Tris gel in MOPS buffer (ThermoFisher Scientific) and transferred to nitrocellulose membranes (GE Healtcare Life Sciences). The membranes were blocked with 5% (w/v) milk in PBS buffer supplemented with 0.1% Tween 20 for 1h at room temperature. The membranes were treated with primary antibody (3E8, 1:25 dilution) in 0.25 % (w/v) milk in PBS supplemented with 0.1 % Tween 20 and incubated at room temperature for 1 h. After several rounds of washing with PBS buffer supplemented with 0.1 % Tween 20, the membranes were incubated with HRP-conjugated anti-rat secondary antibody (1:5000 dilution, Sigma-Aldrich A9037) in 0.1 % (w/v) milk in PBS supplemented with 0.1 % Tween 20 and incubated at room temperature for 1 h. Antibodies were detected with SuperSignal West Pico Chemiluminescent Substrate (ThermoFisher) and the membranes were scanned with ChemoCam Advanced Fluorescence imaging system (Intas Science Imaging).

### Mass spectrometric identification of crosslinking sites

Sample for crosslinking was prepared by mixing core TFIIH, XPA, XPG and bifurcated DNA scaffold (see cryo-EM sample preparation) in 1:2:2:1.5 molar ratio in a final buffer containing 150 mM KCl, 10% glycerol, 2 mM MgCl_2_, 20 mM KOH:HEPES pH 7.5 and 5 mM β-mercaptoethanol. The reaction was incubated for 20 min at room temperature before applying to the Superose 6 increase 3.2/300 column equilibrated in final buffer used for the complex formation. Fractions were analyzed by SDS-PAGE and Coomassie staining. Appropriate fractions were pooled, supplemented with 2 mM BS3 crosslinker (ThermoFisher Scientific) and incubated for 30 min at 30 °C. Reactions were quenched with ammonium bicarbonate (200 mM final) and further incubated for 10 min at 30 °C.

Crosslinked proteins were reduced with 10mM dithiothreitol (DTT) for 30min at 37 °C, followed by alkylation with 40mM iodoacetamide for 30 min at 25 °C. The proteins were digested overnight at 37 °C in the presence of 1M urea with trypsin in 1:20 (w/w) enzyme-to-protein ratio. The samples were acidified with formic acid (FA) to 0.1% (v/v) final concentration and acetonitrile (ACN) was added to 5% (v/v) final concentration. Peptides were bound to Sep-Pak C18 50 mg sorbent cartlidge (Waters), washed with 5% ACN, 0.1% FA (v/v) and eluted with 80% ACN, 0.1%FA (v/v), dried under vacuum and resuspended in 30 µl 30% ACN, 0.1% trifluoroacetic acid (TFA) (v/v). The sample was fractionated on Superdex Peptide PC3.2/30 column (GE Healthcare) at a flow rate of 50 µl/min of 30% ACN, 0.1% (v/v) and 100 µl fractions corresponding to elution volume 0.9ml-2.2ml were collected, dried under vacuum and resuspended in 20 µl 2% ACN, 0.05% TFA (v/v).

Crosslinked peptides were analyzed on Orbitrap Fusion Tribrid Mass Spectrometer (Thermo Scientific), coupled to Dionex UltiMate 3000 UHPLC system (Thermo Scientific) equipped with an in-house-packed C18 column (ReproSil-Pur 120 C18-AQ, 1.9 µm pore size, 75 µm inner diameter, 30 cm length, Dr. Maisch GmbH). Samples were analyzed as 5µl injections, separated on 118 min gradient: mobile phase A - 0.1% FA (v/v); mobile phase B - 80% ACN, 0.08% FA (v/v). The gradient started at 8% B and increased to 48% B in 106 min, then keeping B constant at 90% for 6 min, followed by reequilibration of the column with 5% B. The flow rate was set to 300 nl/min. MS1 spectra were acquired with the following settings: resolution - 120,000; mass analyzer - Orbitrap; mass range - 380- 1500 m/z; injection time - 60ms; automatic gain control target - 4 × 105. Dynamic exclusion was set to 9 s. For samples of elution volume up to 1.7ml only charges 3-8 were included. For subsequent samples – charges 2-8 were included. MS2 spectra settings: resolution - 30,000; mass analyzer – Orbitrap; injection time - 128 ms; automatic gain control target - 5 × 104; isolation window - 1.6 m/z. Fragmentation was enforced by higher-energy collisional dissociation and varied for the three injections at 30%, 28% and stepped collision energy with step +/- 2% at 28%.

Raw files were converted to mgf format with ProteomeDiscoverer 2.1.0.81 (Thermo Scientific): signal-to-noise ratio - 1.5; precursor mass - 350–7,000 Da. For identification of crosslinked peptides, files were analysed by pLink (v. 1.23) (pFind group)^43^ with the following settings : crosslinker – BS3; digestion enzyme – trypsin; missed cleavage sites – 2; fixed modification - carbamidomethylation of cysteines; variable modification - oxidation of methionines; precursor mass tolerance filtering – 10 ppm; fragment ion mass tolerance - 20 ppm; false discovery rate – 1%. Cross-link files were also identified with pLink (v. 2.3): linkers – BS3; missed cleavages – 3; precursor tolerance – 6ppm; fragment tolerance – 20ppm; peptide length – 4-60; filter tolerance – 10 ppm; spectral level false discovery rate – 1%. The sequence database contained all proteins within the complex. Cross-linking figures were created with XiNet^44^ and Xlink Analyzer plugin in Chimera^45^.

### Cryo-EM sample preparation and image processing

Sample was prepared by mixing pre-annealed DNA scaffold (5’- CAAAGTCACGACCTAGACACTGCGAGCTCGAATTCACTGGAGTGACCTC -3’; 5’- GAGGTCACTCCAGTGAATTCGAGCTCGCAACAATGAGCACATACCTAGT - 3’) with core TFIIH, XPA and XPG:E791A in 1.5:1:3:3 molar ratio in final buffer containing 150 mM KCl, 20 mM KOH:HEPES pH 7, 10 % glycerol, 2 mM MgCl_2_ and 5 mM β-mercaptoethanol. XPG:E791A endonuclease mutant was used to prevent DNA cleavage during the sample preparation^46^. The sample mixture was applied to a sucrose gradient in order to purify the complex from excess factors and fix it with glutaraldehyde^47^. The sucrose gradient was prepared with BioComp Gradient Master 108 (BioComp Instruments) by mixing equal volume of heavy (30 % (w/v) sucrose, 150 mM KCl, 2 mM MgCl_2_, 5 mM β-mercaptoethanol, 20 mM KOH-HEPES pH 7.5 and 0.1 % glutaraldehyde) and light solutions (10 % (w/v) sucrose, 150 mM KCl, 2 mM MgCl_2_, 5 mM β- mercaptoethanol and 20 mM KOH-HEPES pH 7.5) in 5 ml centrifugation tubes. After 16 h of centrifugation at 4°C and 175, 000 xg the gradient was fractionated and glutaraldehyde was quenched with lysine (50 mM final) and aspartate (20 mM final). Fractions were dialysed in Slide-A-Lyzer MINI Dialysis Devices (2ml and 20 kDa cut-off) (ThermoFisher Scientific) for 10 h against buffer containing 100 mM KCl, 20 mM KOH-HEPES pH 7.5, 1 mM MgCl_2_, 1 mM DTT, 0.5% glycerol (v/v) and 0.004% n-octyl glucoside (w/v). Dialysed samples were immediately used for cryo-grid preparation. 4 ul of sample was applied to glow-discharged R2/2 gold grids (Quantifoil) which were blotted for 5 s and plunge-frozen in liquid ethane with a Vitrobot Mark IV (FEI) operated at 4 °C and 100 % humidity.

Micrographs of the sample were acquired on a FEI Titan Krios G2 transmission electron microscope with a K2 summit direct electron detector (Gatan). Data acquisition was automated with FEI EPU software package. Micrographs were acquired at a nominal magnification of 130,000x (1.05 Å per pixel) using a dose rate of 4.55 e^-^ per Å^2^ per s over the time of 9 s which resulted in a total dose of 41 e^-^ per Å^2^ fractionated over 40 frames. CTF correction, motion correction and particle picking was done on-the-fly using Warp^48^. Automated picking in retrained BoxNet implemented in Warp^48^ yielded a total of 1,354,997 particles from 8,993 micrographs, which were further subjected to 2D classification in CryoSPARC^49^. After 2D cleaning, 950,000 particles were used for heterogeneous refinement in CryoSPARC. Three ab-initio classes obtained from the first 300,000 particles picked during data acquisition were used as an input for the refinement. The class showing clear core TFIIH features was further 3D classified into 6 classes using RELION-3^50^. Particles corresponding to the best 3D class were subjected to CTF refinement and Bayesian polishing. Particles were 3D refined and post-processed with automatic B-factor determination in RELION. Final map showed an overall resolution of 3.6 Å according to the gold-standard Fourier shell correlation (FSC) 0.143 criterion with an applied B-factor of −110.02 Å^-2^. Due to flexibility of peripheral regions of core TFIIH, we improved the map quality for 5 different regions of the complex by focused 3D classification and refinement (processing tree is depicted in Extended Data Fig. 2d). The classifications were performed with particles contributing to the final map without image alignment to speed up the calculations. Masks encompassing the regions of interest were created with UCSF Chimera^35^ and RELION. 3D classification of the DNA duplex revealed two alternative DNA conformations within the complex (Extended Data Fig. 2d).

### Model building

The final cryo-EM map and focused classified maps were used for model building. The final map was denoised in Warp 1.0.6^48^. Structures of ATPase lobe 1 and 2 of XPB, XPD, p44 vWA-like domain and p52 C-terminus (residues 383-458) from the TFIIH structure (PDB code 5OF4)^6^, as well as the crystal structure of p34 vWA-like domain bound to p44 RING domain (PDB code 5O85)^51^ were rigid-body fitted into our cryo-EM density in UCSF Chimera^35^ and manually adjusted in COOT^52^. Due to high quality of the EM density, the NTE domain and part of the DRD domain (residues 71-199 and 266-300), as well as the p52 region that interacts with XPB (residues 290-382) were built *de novo* guided by secondary structure prediction in PSIPRED^53^ and bulky amino acid side chains as sequence registers. In case of XPD we did not observe EM density corresponding to residues 273- 325, so we removed this part of the structure. We observed a very strong density for the iron-sulfur cluster indicating that the ligand was not damaged or dissociated during protein expression and purification, as well as sample preparation for cryo-EM. The N-terminal region of p52 (residues 18-289) and zinc-fingers belonging to subunits p34 and p44 were modelled with SWISS-Model^54,55^ based on the yeast p52 counterpart (PDB code 5OQJ)^7^ and manually adjusted in COOT. Interestingly, the p34 zinc finger region in human contains additional cysteine (C257) and histidine (H258) residues not present in the yeast counterpart which allows binding of an additional zinc ion. The smallest TFIIH subunit TTDA (p8) was generated in Modeller^56^ with the yeast TTDA structure as a reference (PDB code 5OQJ)^7^, rigid-body fitted in our density using UCSF Chimera and manually adjusted in COOT. The NMR structure of truncated human XPA (PDB code 1XPA)^57^ was also docked in our density as a rigid body and adjusted in COOT. We observed additional helical density that extends from the C-terminus of the docked structure towards the ATPase lobe 2 of XPB when the map is filtered to lower resolution. Secondary structure prediction with PSIPRED shows that the 22 residues that follow docked XPA C-terminus form a helix, so we extended the C-terminal helix in COOT guided by the cryo-EM density (Extended Data Fig. 3h).

DNA sequence was assigned based on the position of the DNA duplex-single strand junction, however protein binding to the junction could induce additional DNA melting so register shifts cannot be excluded. DNA duplex was built by docking ideal B-DNA into the density, followed by manual adjustments in COOT. Several rounds of real space refinement and geometry optimisation with secondary structure restraints (including base pairing and base stacking restrains) were performed in PHENIX^58^. The DNA duplex-single strand junction and single-strand extensions were manually built in COOT. The EM density for the 5’-3’ DNA single strand showed clear separation of sugar, phosphate and DNA bases for nucleotides A30-G36 and for C40-A41. The decreased quality of EM map for nucleotides G37-A39 and T42, presumably due to increased flexibility of DNA between XPD helicase lobes, allowed the trajectory of DNA to be determined, but the nucleotides were positioned manually in COOT guided by the structure of NS3 helicase in complex with DNA^59^ and real space refined in PHENIX. All core TFIIH subunits, XPA and DNA were first real-space refined in PHENIX separately in their corresponding focused classified maps. Then, all components were combined and real-space refined together in the global map. The final model was validated using Molprobity^60^.

### Data availability statement

The coordinates for the TFIIH-XPA-DNA complex are available in the PDB (code XXXX), and the cryo-EM maps are available in EMDB (code YYYY).

## EXTENDED DATA FIGURE LEGENDS

**Extended Data Figure 1.**
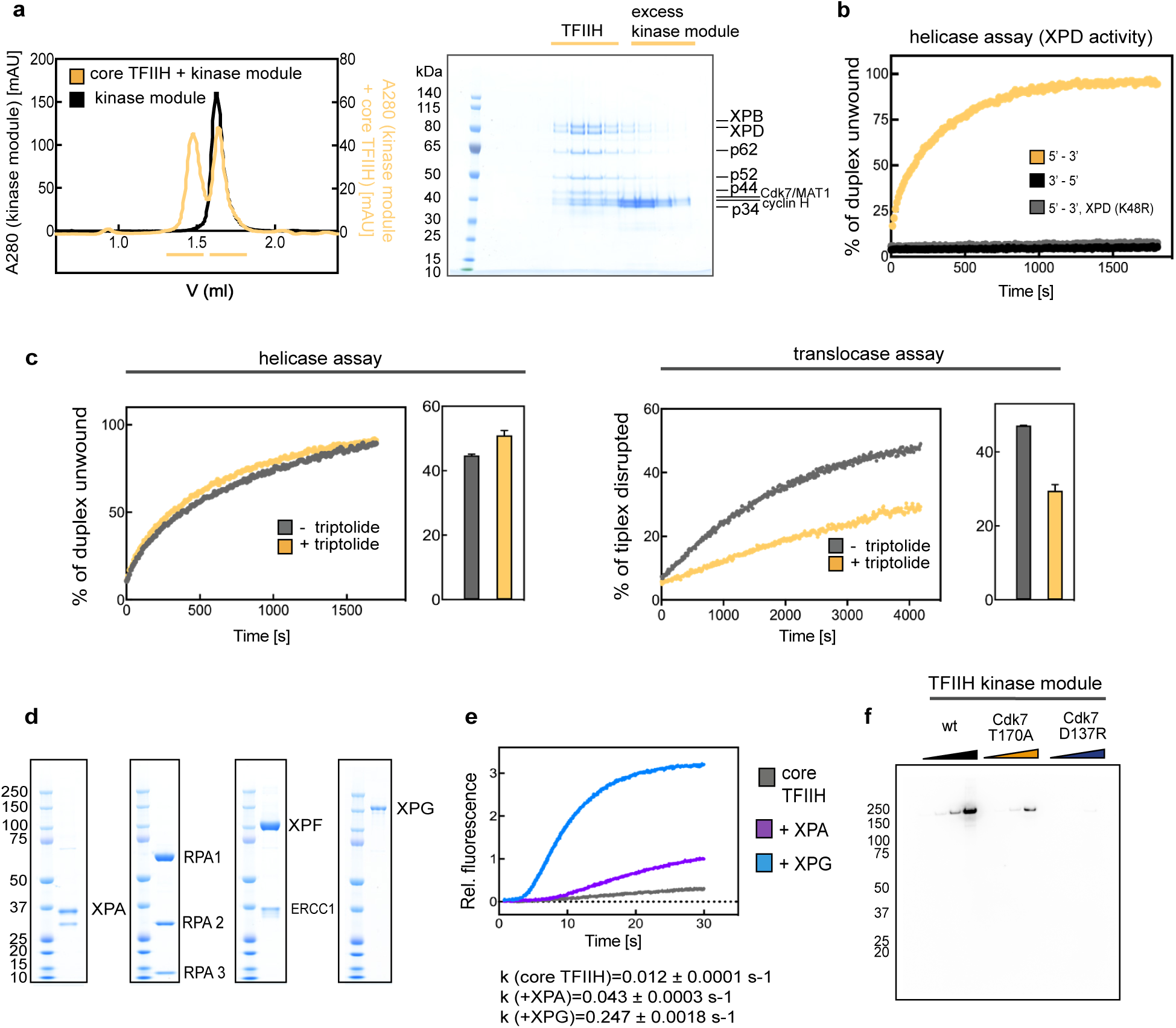
Biochemical characterization of TFIIH helicases. **a.** Reconstitution of the TFIIH complex by size-exclusion chromatography. Chromatograms show elution profile for kinase module in black and the mixture of the core TFIIH and the kinase module in yellow. Two peaks detected for the mixture of the core TFIIH and the kinase (yellow lines) were resolved by SDS-PAGE. First peak contains reconstituted TFIIH and the second peak contains the excess kinase module. Gels also demonstrate the purity of TFIIH preparations. **b.** Polarity of XPD helicase. DNA unwinding was monitored by a fluorescence resonance energy transfer - based assay. Fluorescence traces of the core TFIIH unwinding in the 5’ – 3’ and the 3’ – 5’ direction are shown in yellow and black, respectively. DNA unwinding of the core TFIIH containing XPD K48R mutant is shown in gray. Shown traces are representative of 3 independent experiments. **c.** DNA translocation by the core TFIIH is mediated by XPB. DNA unwinding and triplex disruption were monitored by a fluorescence resonance energy transfer - based assay. Helicase (left) and translocase (right) activities of the core TFIIH in the presence (yellow) or absence (gray) of triptolide (100 µM final) are shown. Triptolide specifically inhibits the XPB enzyme^10^ and affects the core TFIIH translocation, without affecting the XPD helicase activity, indicating that XPB mediates DNA translocation. Bar graphs show the percentage of unwound duplex after 300 s (left) and the percentage of disrupted triplex after 4000 s (right) for 2 independent replicates. Error bars represent s.d. from the mean values. **d.** Purified NER factors resolved by SDS-PAGE and visualized by Coomassie staining. **e.** Stopped-flow measurement of DNA unwinding. TFIIH core (gray) was pre-incubated with excess XPA (purple) or XPG (blue) and rapidly mixed with ATP in a stopped-flow apparatus. DNA unwinding was monitored by a fluorescence resonance energy transfer - based assay. Fluorescence traces represent the average of 5 measurements. Initial linear parts of the traces were fitted with Prism software to obtain the initial rate of DNA unwinding, as indicated below the graph. **f.** Activity of kinase module variants. Different kinase module variants were incubated with the dephosphorylated yeast RNA-polymerase II and the phosphorylation status of the C-terminal domain of Rpb1 was probed by Western blotting. CDK7:T170A exhibits a much weaker kinase activity compared to the wild type CDK7, while CDK7:D137R mutant is kinase dead.

**Extended Data Figure 2.**
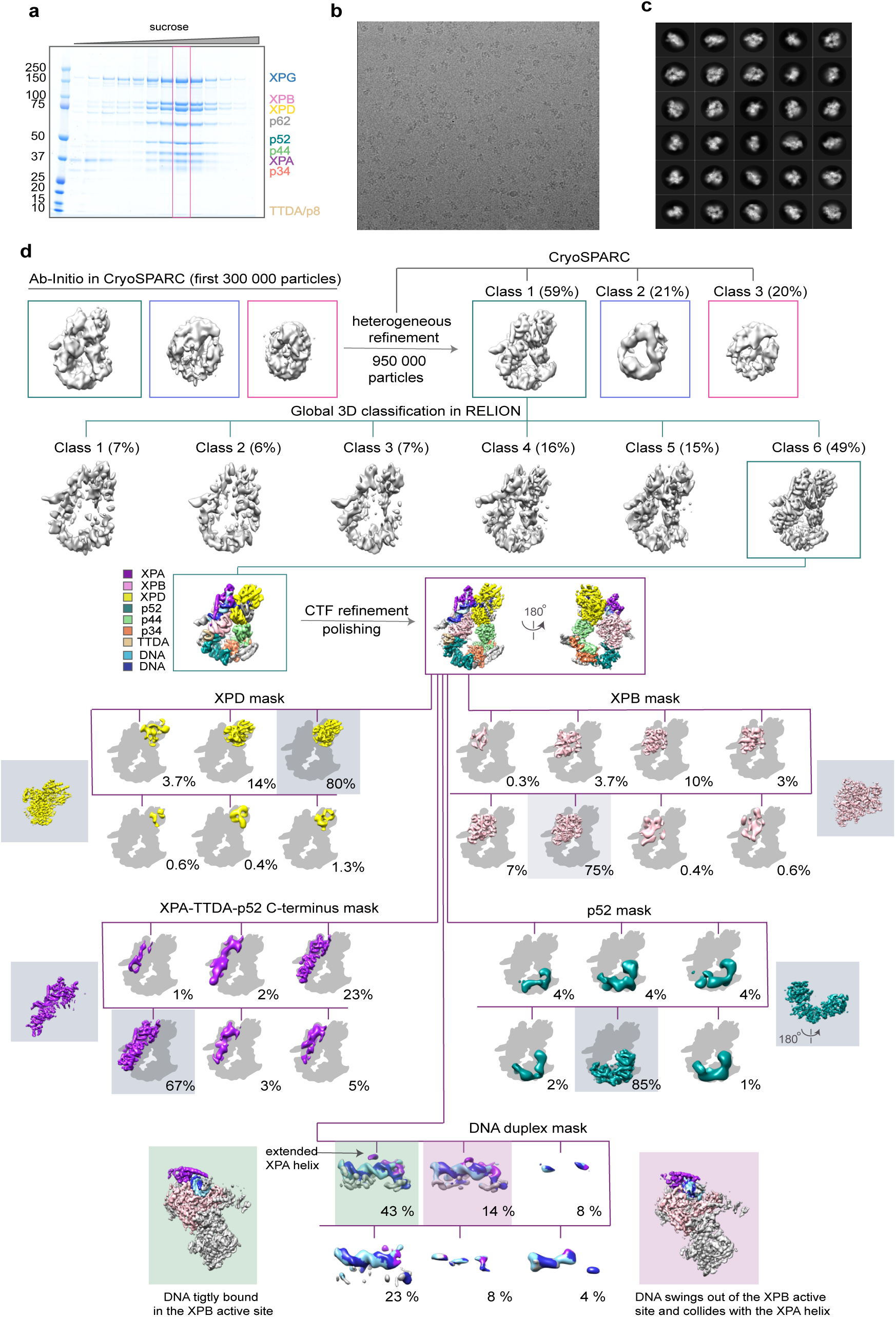
Cryo-EM data processing. **a.** Core TFIIH-XPA-XPG-DNA complex formation for cryo-EM. Sucrose gradient fractions from the control without glutaraldehyde were analyzed by SDS-PAGE and Coomassie staining. Framed fraction in the presence of glutaraldehyde was used for structure determination. **b.** Representative micrograph shows nicely spaced individual particles. **c.** Thirty representative 2D classes obtained after the final 3D classification. **d.** Processing tree for structure determination of the core TFIIH-XPA-DNA complex. Particle class-distribution after 3D classification is indicated next to the corresponding map. The final map obtained by global refinement is color coded by subunits as indicated. 5 focused classifications used to improve different parts of the structure are shown over the gray TFIIH silhouette for context. Classes used for the refinement are indicated with colored background and the refined structure is shown next to the corresponding 3D classes.

**Extended Data Figure 3.**
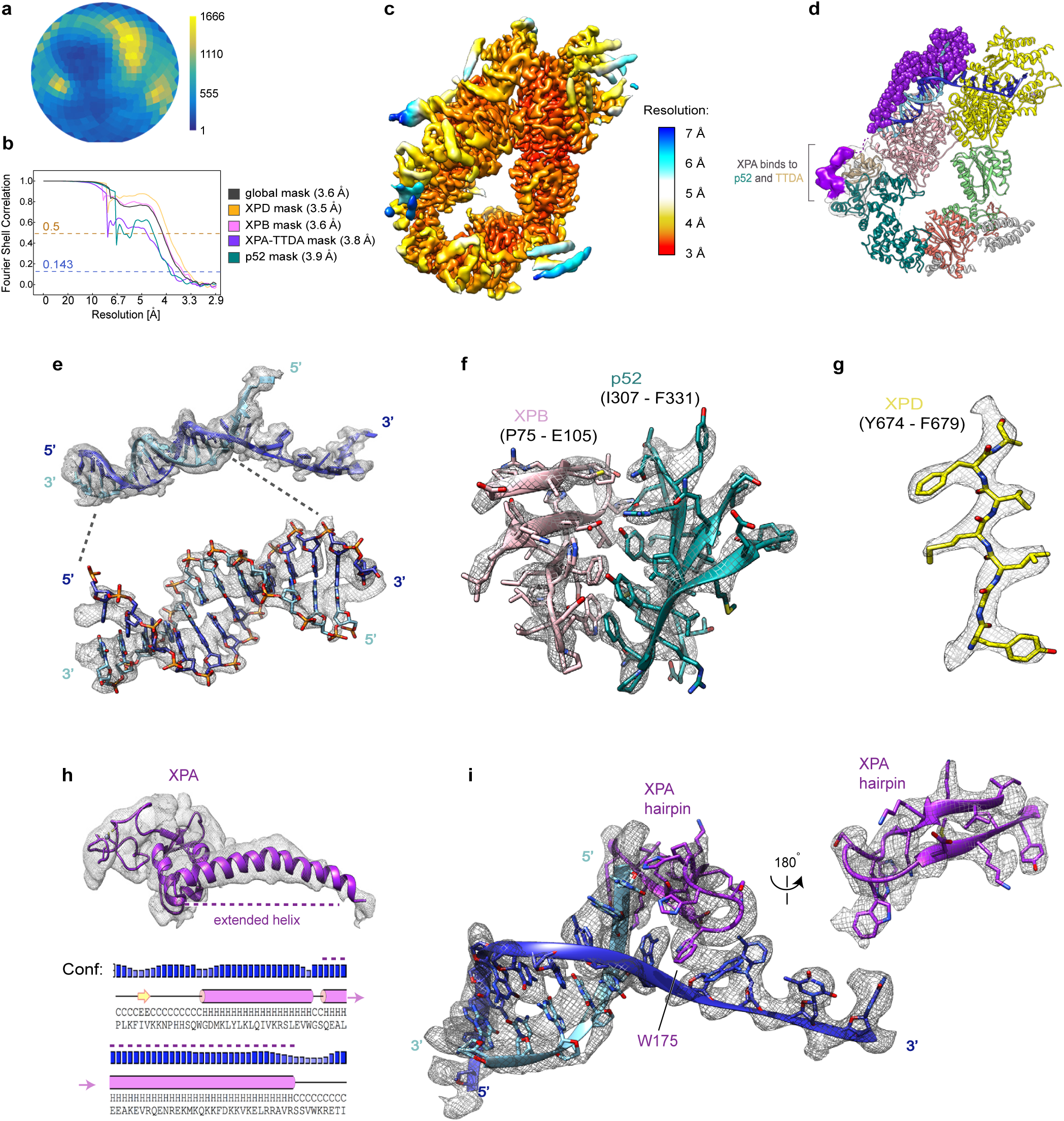
Cryo-EM map quality. **a.** Angular distribution plot for particles contributing to the final reconstruction. Number of particles assigned to a particular orientation is color-coded as indicated. **b.** Fourier shell correlation plots for the final and focused-classified maps. **c.** Local resolution estimates for the final map. **d-i.** Examples of the final map quality. d. Additional XPA density (purple) contacting p52 dimerization domain and TTDA/p8 is observed at lower resolution and it was not used for model building. XPA is shown as purple spheres and TFIIH subunits are color coded as in Fig. 2a. e. Cryo-EM density corresponding to the bifurcated DNA scaffold. Close-up view of the double stranded DNA region shows a clear separation of the density corresponding to phosphates, sugar and DNA bases. f. De novo built interface between the XPB and p52 subunits. Bulky side chains were used to assign the protein register. g. Individual beta-strand from the second helicase lobe of XPD. The high map quality allows the protein backbone tracing and the placement of side-chains. h. Cryo-EM density for the extended XPA helix directs the placement of the secondary structure element in the lower resolution map. PSIPRED tool^53^ predicts the formation of the long helix with high confidence. i. Cryo-EM density for the XPA intercalating hairpin. W175 is indicated.

**Extended Data Figure 4.**
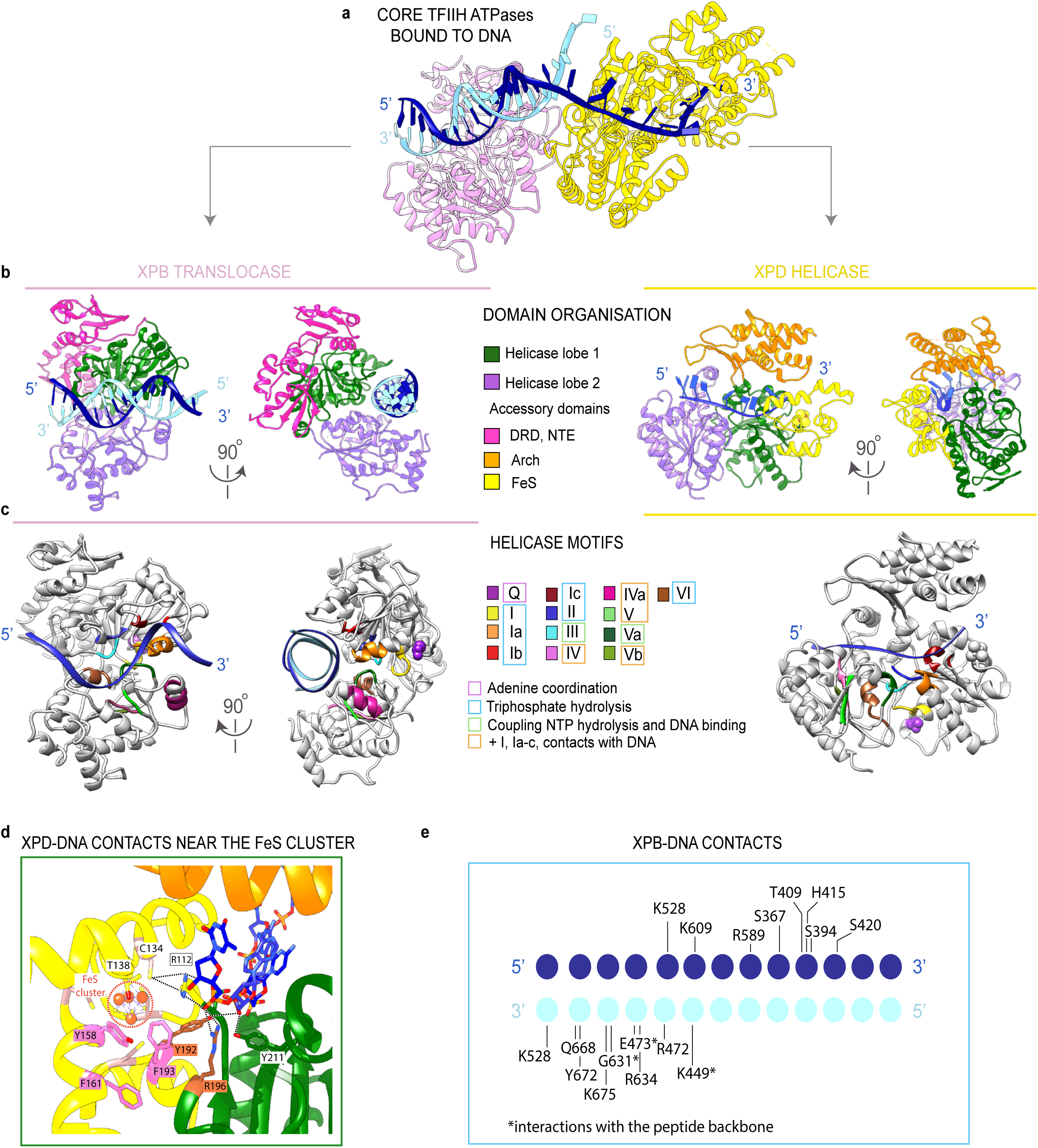
Detailed analysis of XPB and XPD helicase structures. **a.** Ribbon representation of XPB (pink) and XPD (yellow) bound to the DNA. **b.** Domain organization of XPB (left) and XPD (right). Helicase lobes and accessory domains are color coded as indicated. **c.** Helicase motifs colored for XPB (left) and XPD (right) according to^61^. In XPB structure it is clearly visible that the 5’-3’ DNA strand interacts with the helicase motifs. **d.** XPD-DNA interactions near the FeS cluster. The FeS domain is shown in yellow, the Arch domain in orange and the ATPase lobe 1 in green. Residues implicated in lesion scanning are highlighted in brown and aromatic residues surrounding FeS cluster in pink. **e.** Schematic representation of XPB-DNA interactions.

**Extended Data Figure 5.**
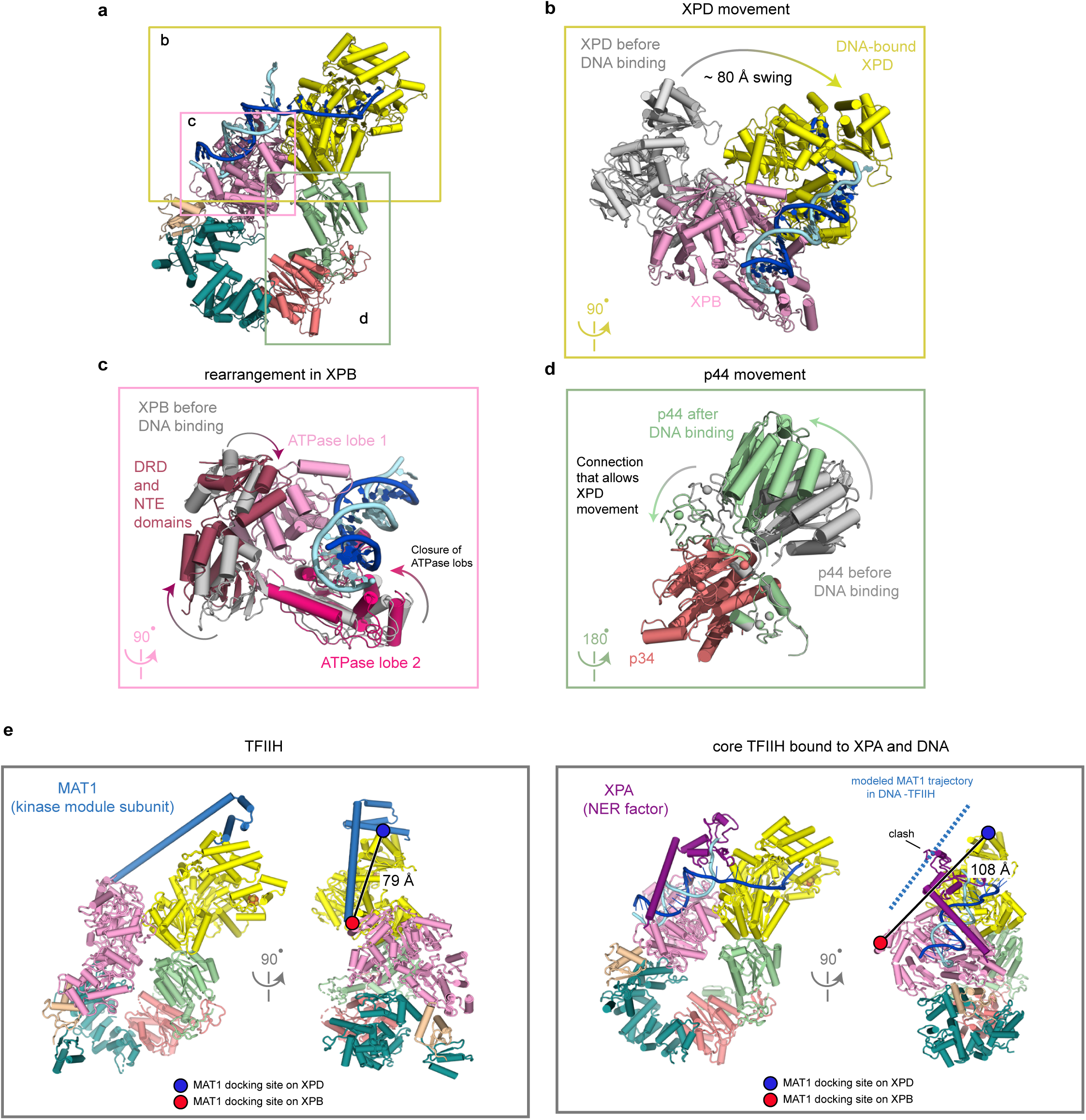
Large-scale structural changes in TFIIH. **a.** The core TFIIH-DNA structure shown for context and color-coded as in Fig. 2a. **b-d.** Superpositions of different TFIIH subunits before and after DNA binding show inter- and intra-subunit flexibility. TFIIH structure in absence of DNA is always shown in gray. The structure was modeled by fitting TFIIH domains built here into the EM density for the kinase-bound TFIIH^6^ for easier comparison, followed by a real-space refinement in PHENIX. b. The structures with and without the DNA were aligned on the ATPase lobe 1 of XPB. XPD swings ~80 Å during DNA loading (R272 in corresponding structures was used for the distance measurement). c. Internal rearrangements in XPB. XPB domains are color-coded as follows: NTE and DRD domains in dark pink, ATPase lobe 1 in pink and ATPase lobe 2 in hotpink. Binding to DNA duplex induces closure of XPB ATPase lobes and substantial rearrangement of accessory domains DRD and NTE. d. Flexible interface between p44 and p34 comprised of several Zn-fingers supports the large-scale movement of XPD during DNA loading. **e.** Structural basis of the core TFIIH activation for DNA repair. Kinase-bound core TFIIH (left) and DNA-bound core TFIIH-XPA (right) structures are shown. Front views show that MAT1 and XPA bridge the core TFIIH ATPases in the kinase-bound structure and in the DNA-bound structure, respectively. Side views were aligned on XPD and show how XPB and XPD reorient during DNA and XPA binding. Distance between the docking sites of MAT1 on XPB (red circle, Q190 residue was used for distance measurement) and XPD (blue circle, V365 was used for distance measurement) on both structures is indicated. The distance increases significantly in the core TFIIH conformation stabilized by DNA and XPA which would induce the kinase module rearrangement on the core TFIIH and may facilitate kinase module dissociation *in vivo*^32^. MAT1 helix trajectory that spans between XPB and XPD was modeled in the DNA-bound structure (dotted blue line) and suggests a potential clash with XPA.

**Extended Data Figure 6.**
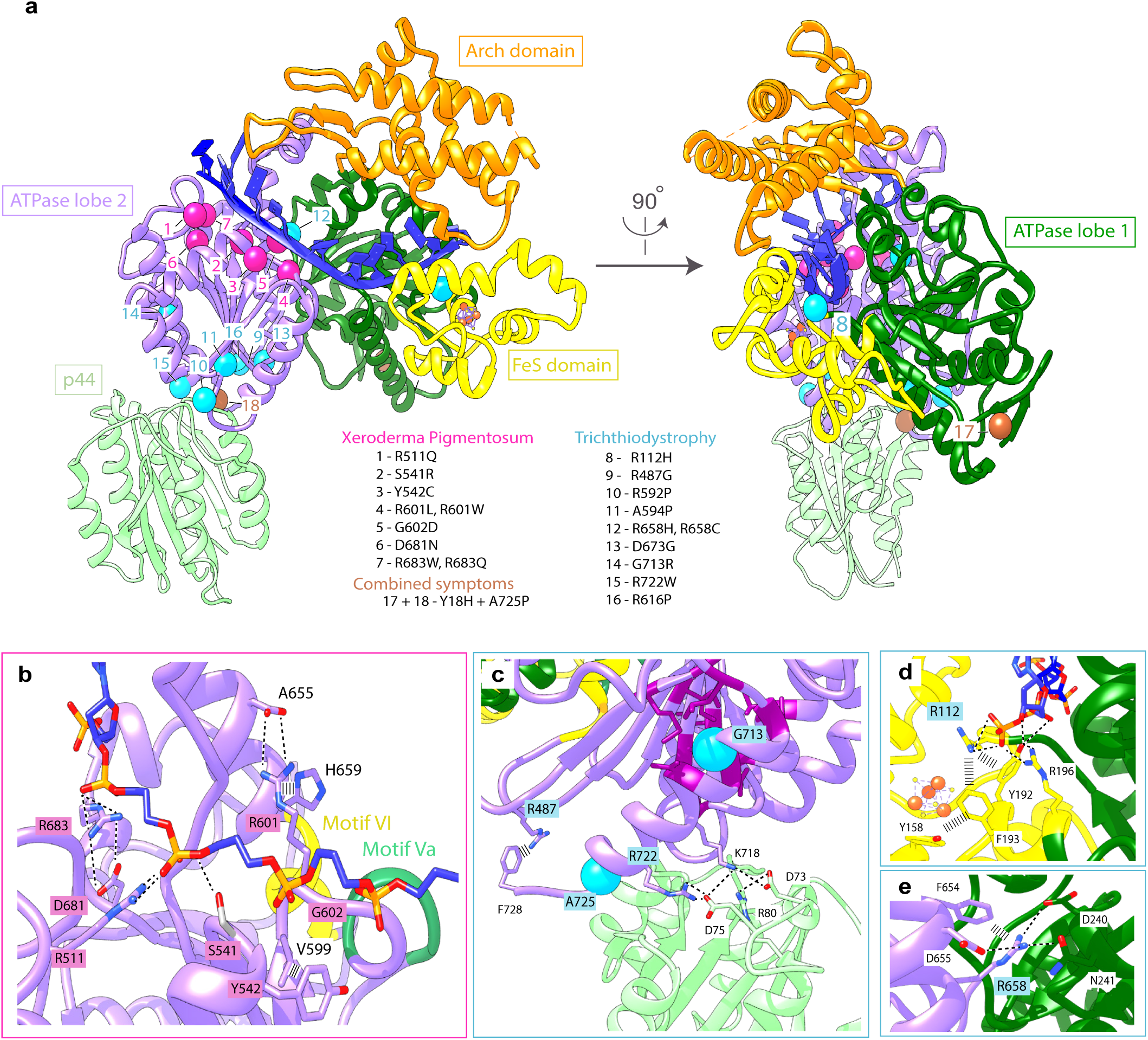
DNA-bound TFIIH core at 3.6Å reveals structural etiology of Xeroderma pigmentosum (XP) and Trichothiodystrophy (TTD) **a.** XP and TTD mutations^3,22,62^ mapped onto XPD-DNA structure are shown as pink and blue spheres, respectively. XP mutations cluster around DNA in the ATPase lobe 2 of XPD and TTD mutations cluster at the XPD-p44 interface. A XP-TTD overlap patient carries 2 mutations^63^, shown as brown spheres. **b.** Zoom-in on XP mutations in the direct vicinity of DNA. Residues mutated in XP are highlighted in pink and their interaction partners in white. Helicase motifs VI and Va are shown in yellow and green, respectively. **c.** XPD-p44 interface. A single helix in XPD is affected by four different TTD mutations. Mutated residues are highlighted in blue. G713 and A725 are depicted as blue spheres. The helix carrying TTD mutations interacts with p44 via two salt bridges, R722 (XPD) - D75 (p44), and K718 (XPD) - D73 (p44). **d.** R112 residue, mutated in TTD, interacts with the DNA backbone and residues surrounding the iron sulfur cluster. **e.** R658 residue, mutated in TTD, engages in the interaction network between the ATPase lobes.

**Extended Data Figure 7.**
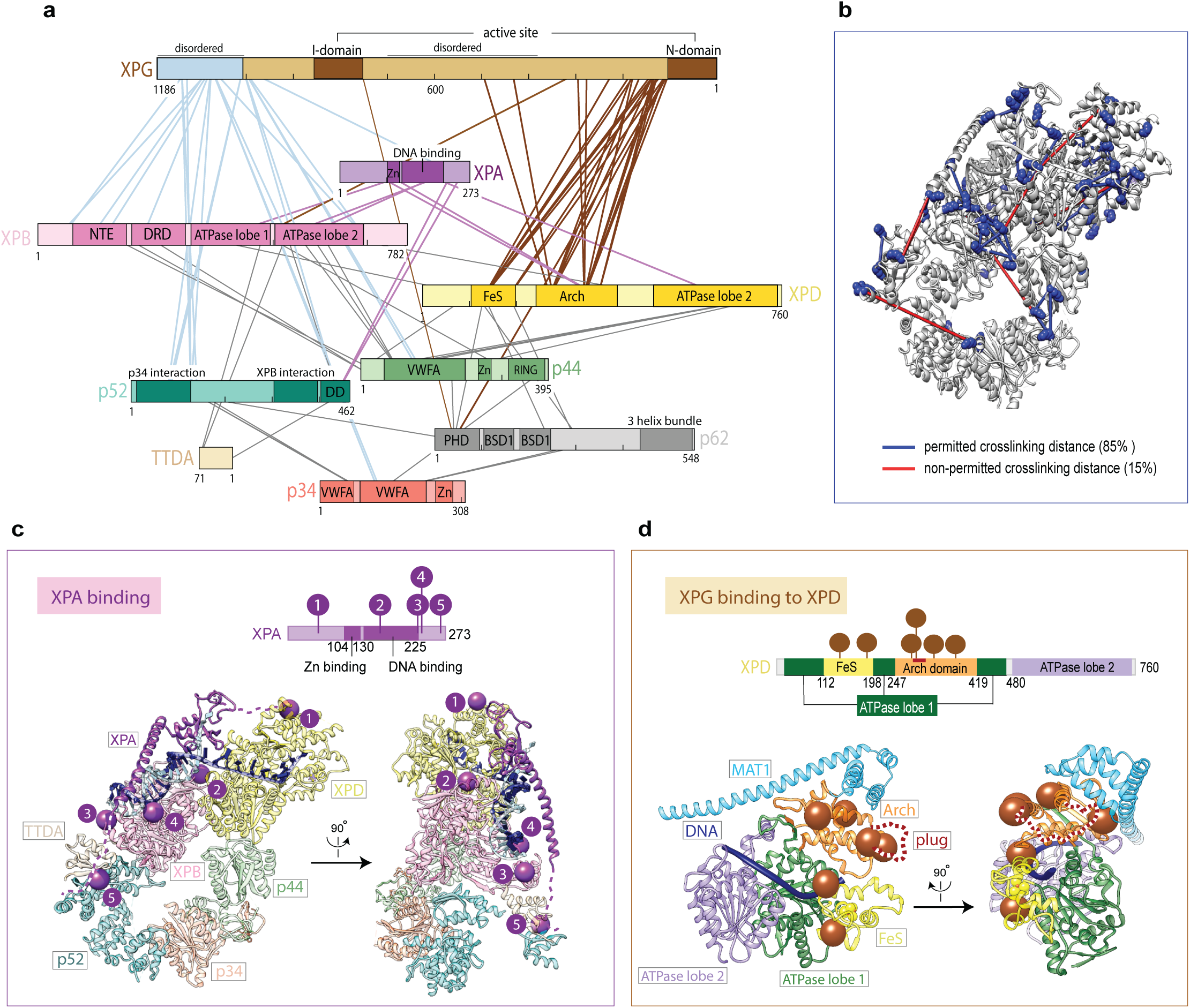
Crosslinking mass-spectrometry network of the core TFIIH-XPA-XPG-DNA complex. **a.** Crosslinks between XPA and core TFIIH are shown in purple, the disordered XPG C-terminus and core TFIIH in blue, and XPG N-terminal region and core TFIIH in brown. Crosslinks between the core TFIIH subunits are in gray. Only crosslinks with the score above 6 are shown. Crosslinks between the subunits not shown here can be found in the Supplementary Table 1. **b.** Validation of crosslinking sites. Crosslinks with the score above 6 were mapped onto core TFIIH-XPA-DNA structure. Colored rods connecting crosslinked residues represent permitted (blue) or non-permitted (red) crosslinking distances, and lysine residues are shown as blue spheres. 85% of mapped crosslink sites fall within the permitted crosslinking distance. **c.** XPA crosslinks mapped onto core TFIIH-XPA-DNA structure. XPA crosslinks are shown as purple spheres. Schematic representation of XPA with mapped crosslinks is shown above the structure. **d.** XPG crosslinks mapped onto XPD-DNA structure. XPD domains are color coded as in Fig. 4, MAT1 subunit of the kinase module is shown in blue and was modeled based on the previous TFIIH structure^6^. XPG crosslinks are shown as brown spheres. Schematic representation of XPD with mapped crosslinks is shown above the structure.

**Extended Data Figure 8.**
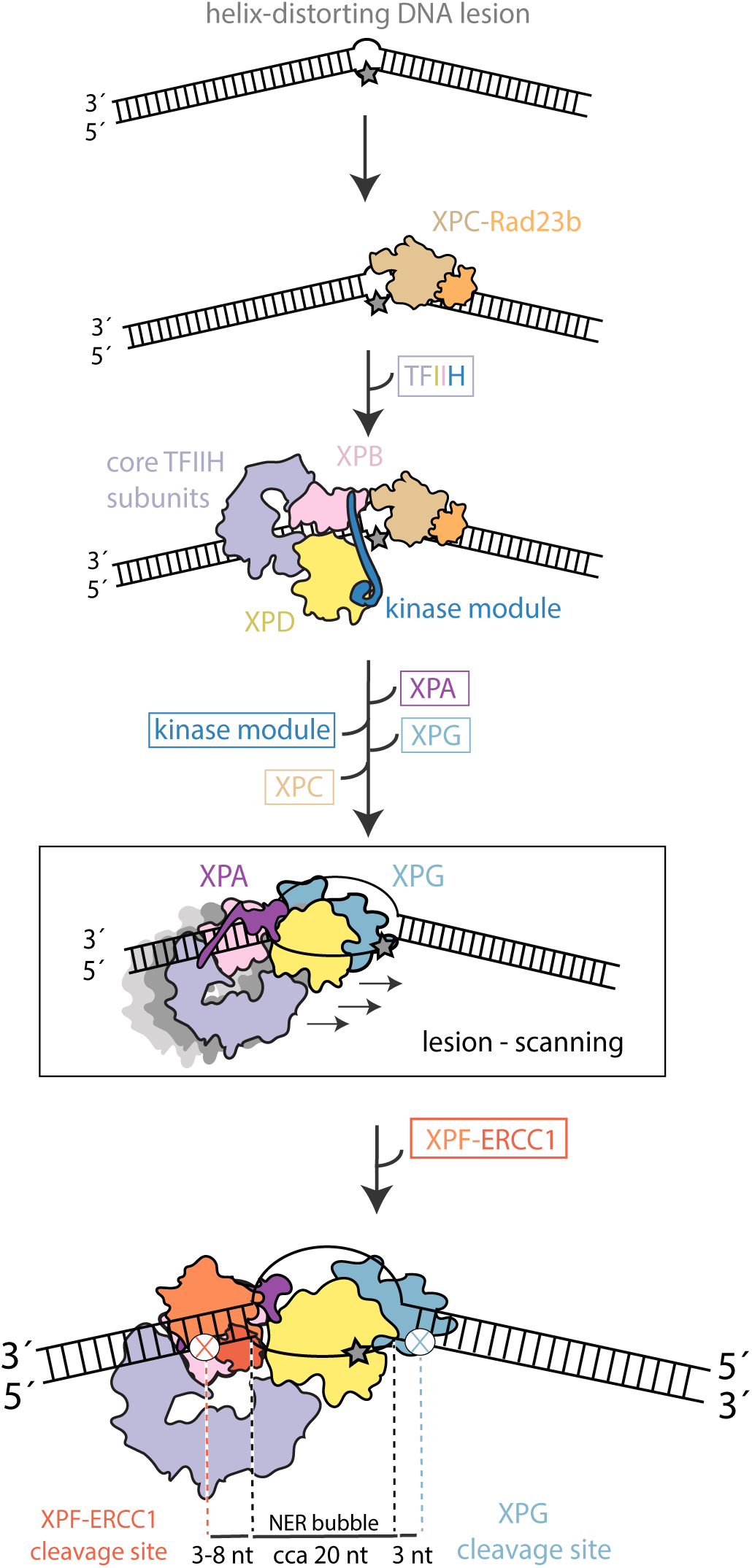
Schematic representation of the NER pathway highlighting new insights. RPA was not included for clarity but it most likely engages the undamaged DNA strand of the repair bubble. Our structure suggests that XPA-XPD footprint on the DNA pre-defines the size of the excised DNA fragment during NER. See main text for more details.

## EXTENDED DATA TABLE

**Extended Data Table 1.**
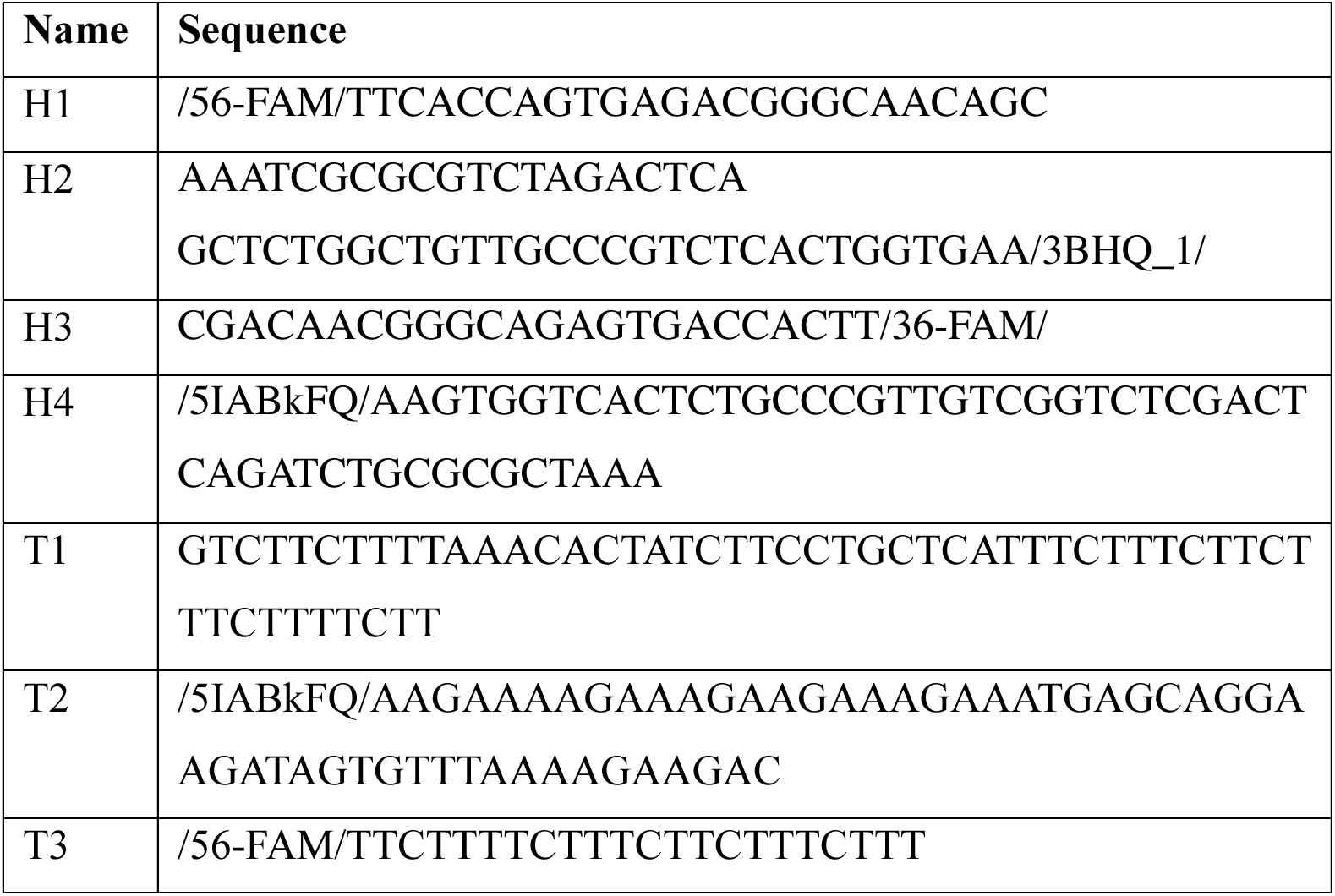
DNA sequences used in this study. All synthetic DNA oligonucleotides were purchased from Integrated DNA Technologies.

## SUPPLEMENTARY INFORMATION

**Supplementary Table 1 Core TFIIH-XPA-XPG-DNA BS3 crosslinking data**

Table of all BS3 crosslinks detected in core TFIIH-XPA-XPG-DNA.

**Supplementary Video 1**

Structural changes in core TFIIH upon activation for DNA repair.

